# Aggressiveness as a latent personality trait of domestic dogs: testing local independence and measurement invariance

**DOI:** 10.1101/117440

**Authors:** Conor Goold, Ruth C. Newberry

## Abstract

Studies of animal personality attempt to uncover underlying or ‘latent’ personality traits that explain broad patterns of behaviour, often by applying latent variable statistical models (e.g. factor analysis) to multivariate data sets. Two integral, but infrequently confirmed, assumptions of latent variable models in animal personality are: i) behavioural variables are independent (i.e. uncorrelated) conditional on the latent personality traits they reflect (*local independence*), and ii) personality traits are associated with behavioural variables in the same way across individuals or groups of individuals (*measurement invariance*). We tested these assumptions using observations of aggression in four age classes (4 - 10 months, 10 months - 3 years, 3 - 6 years, over 6 years) of male and female shelter dogs (N = 4,743) in 11 different contexts. A structural equation model supported the hypothesis of two positively correlated personality traits underlying aggression across contexts: aggressiveness towards people and aggressiveness towards dogs (comparative fit index: 0.96; Tucker-Lewis index: 0.95; root mean square error of approximation: 0.03). Aggression across contexts was moderately repeatable (towards people: intraclass correlation coefficient (ICC) = 0.479; towards dogs: ICC = 0.303).

However, certain contexts related to aggressiveness towards people (but not dogs) shared significant residual relationships unaccounted for by latent levels of aggressiveness.

Furthermore, aggressiveness towards people and dogs in different contexts interacted with sex and age. Thus, sex and age differences in displays of aggression were not simple functions of underlying aggressiveness. Our results illustrate that the robustness of traits in latent variable models must be critically assessed before making conclusions about the effects of, or factors influencing, animal personality. Our findings are of concern because inaccurate ‘aggressive personality’ trait attributions can be costly to dogs, recipients of aggression and society in general.

## Introduction

Studies of non-human animal personality demonstrate that animals show relatively consistent between-individual differences in behaviour, and that the behavioural phenotype is organised hierarchically into broad behavioural dimensions or personality traits (e.g. sociability, aggressiveness or boldness) that further exhibit inter-correlations to form behavioural syndromes (e.g. boldness with aggression; [1–5]). To interpret the complexity inherent in behavioural phenotypes, personality traits and behavioural syndromes are frequently inferred using latent variable statistical models [6], which reduce two or more measured variables (the *manifest* variables) into one or more lower-dimensional variables (the *latent* variables), following work in human psychology [7–10].

Many animal personality studies use *formative* models, such as principal components analysis, that construct composite variables comprised of linear combinations of manifest variables. However, formative models impose only weak assumptions about the relationships between latent variables and manifest variables [6,11]. For instance, formative models do not require manifest variables to be correlated with one another or illustrate internal consistency [11]. Because behavioural variables comprising personality traits are expected to correlate with each other [4], the utility of formative models to revealing underlying personality traits has been criticised in both animals [12,13] and humans [10,11,14,15]. Instead, researchers are increasingly using reflective models, such as factor analysis, including confirmatory approaches such as structural equation modelling (see [1,16–18]). Reflective models regress measured behaviours on one or more latent variables, incorporating measurement error and possibilities to compare a priori competing hypotheses [1,16,19].

Whilst reflective models offer a powerful framework to examine the latent variable structure of animal behaviour [19], they impose certain assumptions on the interpretation and modelling of latent variables that have received scrutiny in human psychology but are rarely discussed in studies of animal personality. Two foundational assumptions are *local independence and measurement invariance*. Local independence implies that manifest variables should be independent of each other conditional on the latent variables [20,21]. For example, given a continuous latent variable *θ* (e.g. boldness) and two binary manifest variables *Y*_1_ and *Y*_2_ that can take the values 0 and 1, the item response theory model asserts that *P*(*Y*_1_ = 1, *Y*_2_ = 1|θ) = *P*(*Y*_1_ = 1|θ)*P*(*Y*_2_ = 1|θ). As such, the latent variables should ‘screen off’ any covariance between manifest variables. Measurement invariance implies that the latent variables function the same (i.e. are invariant or equivalent) in different subsets of a population or in the same individuals through time [21–25]. In the previous example, this means that the expected values of the manifest variables *Y*_1_ and *Y*_2_ should remain the same across different groups, ΰ (e.g. sex or different populations), for any fixed value of the latent variable *θ_x_* e.g. *E* (*Y* _1_*θ*_x_ = *E* (*Y*_1_ *θ_x_*, *ΰ*). For studies of personality, violations of local independence or measurement invariance highlight instances where the personality traits do not completely explain variation in the manifest variables, which may lead to misleading conclusions about the differences between individuals as a function of trait scores [25–27].

The goal of this study was evaluate local independence and measurement invariance in behavioural data on domestic dogs (*Canis lupus familiaris*). Dog personality has been of scientific interest for decades [28–30], both to predict the behaviour of dogs at future time points [31] and to elucidate behavioural traits pertinent to dogs’ domestication history [32–35]. Research on personality in dogs has led to different numbers and composition of hypothesised personality traits with little consensus on how such traits should be compared within and between studies [36–38]. Dog personality studies frequently involve collection of data on a wide range of behaviours and, as a result, latent variable models are popular to reduce behavioural data into personality traits or dimensions [29]. Importantly, the predictive value of personality assessments in dogs has been inconsistent [31,39–43], perhaps most prominently in shelter dog personality assessments (e.g. see [31] for a review). Assessments of aggression are of particular concern, where aggression has been divided into different aggressiveness traits, such as owner-, stranger-, dog- or animal-directed factors [29,37,44,45]. Improving inference about aggressiveness in dogs is important because dog bites are a serious public health concern [46], especially for animal shelters rehoming dogs to new owners, and aggressive behaviour is undesirable to many organisations using dogs for various working roles [47].

Evaluating local independence and measurement invariance could help refine applied personality assessments on dogs. Local independence may be violated in standardised test batteries (a common assessment method; [48–50]) because the sequential administration of different behavioural subtests means that how dogs responds to one sub-test may influence their subsequent behavioural responses, as well as the responses of the dog handlers [31]. Identifying local independence could, thus, highlight which sub-tests can be interpreted as providing independent information. Local independence is also relevant to the development and analysis of dog personality questionnaires completed by dog owners, because the order in which the questions are presented or redundancy in the content of questions can lead to dependencies between participant responses not explained by the questionnaire’s intended focus on the dog’s behaviour [51].

Scientists are also concerned with understanding personality differences in dogs across a variety of conditions, including ontogeny, age, sex, breed and neuter status (e.g. [37, 42, 52–54]). Evaluating measurement invariance in personality assessments would allow researchers to confirm whether differences between individuals or groups of individuals in personality assessments reflect credible differences in personality trait scores or whether additional, unaccounted for factors are driving the differences. While it may be unrealistic for measurement invariance to hold in all instances, it is important to establish whether it holds for personality traits across basic biological variables such as age and sex, which are generally applicable to dog populations undergoing personality assessment and have previously been found to show interactions with personality traits, including playfulness, sociability, curiosity and aggressiveness [33, 55]. However, apart from van den Berg *et al*. [18] who assessed measurement invariance across breed groups, no studies have confirmed measurement invariance or local independence for personality traits.

In this paper, we assessed local independence and measurement invariance of aggressiveness in shelter dogs using a large sample of data on inter-context aggressive behaviour. First, we decomposed observations of aggression towards people and dogs across contexts into separate aggressiveness traits. Secondly, we assessed whether aggression in different contexts remained associated beyond that explained by latent levels of aggressiveness, testing local independence. Thirdly, we investigated whether the probability of aggression in different contexts assumed to be underpinned by the same aggressiveness trait was measurement invariant with respect to sex and age groups.

## Materials & Methods

### Subjects

Observational data on the occurrence of aggression in 4,743 dogs were gathered from Battersea Dogs and Cats Home’s (UK) observational and longitudinal dog behaviour assessment records (Table 1). The data were from a sample of dogs (N = 4,990) at the shelter’s three rehoming centres during 2014 (including dogs that arrived during 2013 or left in 2015). We selected the records from all dogs that were at least 4 months old, excluding younger dogs because they were more likely to be unvaccinated, more limited in their interactions at the shelter and may have been kennelled in different areas to older dogs. Although dogs were from a variety of heritages (including purebreds and mongrels), the analyses here did not explore breed differences because the accurate visual assessment of breed in dogs with unknown heritage has been refuted [56–58].

**Table 1.** Demographic characteristics of the studied dogs.

| Variable | Mean ± SD / N |
| --- | --- |
| Average age at shelter (years; all ≥ 4 months of age) | 3.75 ± 3.03 |
| Total days at the shelter | 25.13 ± 41.53 |
| Weight (average weight if multiple measurements; kg) | 19.06 ± 10.26 |
| Rehoming centre: London / Old Windsor / Brands Hatch | 2897 / 1280 / 566 |
| Males / females | 2749 / 1994 |
| Neutered<sup>1</sup> before arrival / neutered at shelter / not neutered | 1218 / 1665 / 1502 |
| Relinquished by owners / returned to shelter / strays | 2892 / 260 / 1591 |

<sup>1</sup>358 dogs had unknown neuter status

### Shelter environment

The shelter was composed of three different UK rehoming centres: a high-throughput, urban centre based at Battersea, London with capacity for approximately 150-200 dogs; a semi-rural/rural centre based at Old Windsor with capacity for approximately 100-150 dogs; and a rural centre based at Brands Hatch with capacity for approximately 50 dogs. All dogs arrived in an intake area of their respective rehoming centre and, when considered suitable for adoption, were moved to a ‘rehoming’ area that was partially open to the public between 1000 h and 1600 h. All kennels were indoors. Kennels varied in size, but were usually approximately 4m x 2m and included either a shelf and bedding alcove area, or a more secluded bedding area at the back of the kennel (see [59] for more details). At different times throughout the day, dogs had access to indoor runs behind their kennels. In each kennel block area, dogs were cared for (e.g. fed, exercised, kennel cleaned) by a relatively stable group of staff members, allowing the development of familiarity with staff members and offering some predictability for dogs after arrival at the shelter. Although data on the number of dogs in each kennel were incomplete, in the majority of cases dogs were kennelled singly for safety reasons. The shelter mainly operated between 0800 h and 1700 h each day. All dogs were socialised with staff and/or volunteers each day (often multiple times) except on rare occasions when it was deemed unsafe to handle a dog (when training/behavioural modification proceeded without physical contact). Dogs were provided water ad libitum and fed commercial complete dry and/or wet tinned food twice daily (depending on recommendations by veterinary staff). Dogs received daily tactile, olfactory and/or auditory enrichment/variety (e.g. toys, essential oils, classical music, time in a quiet ‘chill-out’ room).

### Data collection

In the observational assessment procedure, trained shelter employees recorded observations of dog behaviour in a variety of contexts as part of normal shelter procedures. Behavioural observations pertaining to each context were completed using an ethogram specific to that context and recorded in a custom computer system. Multiple observations could be completed each day, although we retained only one observation in each context per day (the least desirable behaviour on that day; see below). The ethogram code that best described a dog’s behaviour in a particular context during an observation was recorded by selecting it from a series of drop-down boxes (one for each context). Although staff could also add additional information in character fields, a full analysis of those comments was beyond the scope of this study. The ethogram for each context represented a scale of behaviours ranging from desirable to undesirable considered by the shelter to be relevant to dog welfare and ease of adoption. Contexts had between 10 and 16 possible behaviours to choose from, some of which overlapped between different contexts. Among the least desirable behaviours in each context was aggression towards either people or dogs (depending on context). Aggression was formally defined as*“Growls, snarls, shows teeth and/or snaps when seeing/meeting other people/dogs*, potentially pulling or lunging towards them”, distinguished from non-aggressive but reactive responses, defined as “*Barks, whines, howls and/or play growls when seeing/meeting other people/dogs, potentially pulling or lunging towards them”*.

Observation contexts included both onsite (at the shelter) and offsite (e.g. out in public parks) settings. For the analyses here, we excluded offsite contexts (which had separate observation categories) because these were less frequently recorded and offsite records were more likely to be completed using second-hand information (e.g. from volunteers taking the dog offsite). We focused on observations of aggression in nine core onsite contexts that were most frequently completed by trained staff members: i) *Handling*, ii) *In kennel*, iii) *Out of kennel*, iv) *Interactions with familiar people*, v) *Interactions with unfamiliar people*, vi) *Eating food*, vii) *Interactions with toys*, viii) *Interactions with female dogs*, ix) *Interactions with male dogs*. For the *In kennel* and *Out of kennel* contexts, recording of aggression towards both people and dogs was possible. If both occurred at the same time, aggression towards people was recorded. Therefore, *In kennel* and *Out of kennel* were each divided to reflect aggression shown towards people and towards dogs only, respectively. This resulted in 11 aggression contexts (Table 2) used as manifest variables in structural equation models to investigate latent aggressiveness traits. The average number of days between successive observations across these contexts and across dogs was 3.27 (SD = 2.08), and dogs had an average of 9.77 (SD = 13.41) observations within each context (N = 416,860 observations in total across dogs, contexts and days). Observations were recorded in the category that best described the scenario. Nonetheless, certain contexts could occur closely in space and time, which were investigated for violations of local independence, as explained below.

**Table 2.** Behavioural observation contexts in which each dog’s reactions were analysed for the presence or absence of aggression.

| Context | Definition |
| --- | --- |
| Handling | Informal handling by people (e.g. stroking non-sensitive areas, touching the collar, fitting a harness or lead). |
| In kennel towards people | People approaching or walking past the kennel. |
| In kennel towards dogs | Dogs in neighbouring kennels or dogs walking past the kennel. |
| Interactions with familiar people | When outside the kennel and familiar people (interacted with at least once before) approach, make eye contact, speak to or attempt to make physical contact with the dog. |
| Interactions with unfamiliar people | When outside the kennel and unfamiliar people (never interacted with before) approach, make eye contact, speak to or attempt to make physical contact with the dog. |
| Out of kennel towards people | When around people outside the kennel who may be a long distance away and who make no attempt to engage with the dog. |
| Out of kennel towards dogs | When around dogs outside the kennel that may be a long distance away and that are not encouraged to interact with the focal dog. |
| Eating food | When eating food (e.g. from a food bowl, or toy filled with food) and people approach within close proximity or attempt to touch the food container. |
| Interactions with toys | When interacting with toys and people approach within close proximity or attempt to touch the toy. |
| Interactions with female dogs | During structured interaction with a female dog, including approaching each other, walking in parallel, and interacting off-lead. Both dogs are aware of each other's presence and are in close enough proximity to engage in a physical interaction. |
| Interactions with male dogs | During structured interaction with a male dog, including approaching each other, walking in parallel, and interacting off-lead. Both dogs are aware of each other's presence and are in close enough proximity to engage in a physical interaction. |

We aggregated behavioural observations across time for each dog into a dichotomous variable indicating whether a dog had or had not shown aggression in a particular context at any time while at the shelter (Table S1). This was performed because the overall prevalence of aggression was low, with only 1.06% of all observations across days involving aggression towards people and 1.13% towards dogs. Thus, the main difference between individuals was whether they had or had not shown aggression in a particular context during their time at the shelter. We interpret aggressiveness here as a between-individual difference variable.

### Validity of behaviour recordings

Validity of the recording of behaviour was assessed separately from the main data collection as part of a wider project investigating the use of the observational assessment method. Ninety-three shelter employees trained in conducting behavioural observations each watched (in groups of 5 – 10 people) 14 videos, approximately 30 seconds each, presenting exemplars of 2 different behaviours from seven contexts (to keep the sessions concise and maximise the number of participants). For each context, behaviours were chosen pseudo-randomly by numbering each behaviour and selecting two numbers using a random number generator. Experienced behaviourists working at the shelter filmed the videos demonstrating the behaviours. Videos were shown to participants once in a pseudo-random order. After each video, participants recorded on a paper answer sheet the behaviour they thought most accurately described the dog’s behaviour based on the ethogram specific to the context depicted. Two of the videos illustrated aggression: one in a combined *Interactions with new* and *familiar people* context (combined because familiarity between specific people and dogs was not universally known) and one in the *In kennel towards dogs* context. The first video had an ethogram of 13 possible behaviours to choose from, and the second had 11 behaviours. The authors were blind to the selection of videos shown and to the video coding sessions with shelter employees.

## Data analysis

All data analysis was conducted in R version 3.3.2 [60].

### Validity of behaviour recordings

The degree to which shelter employees could recognise and correctly record aggressive behaviour from the videos (chosen by experienced behaviourists at the shelter) was determined by the percentage of participants who correctly identified the 2 videos as showing examples of aggression.

### Missing data

Data were missing when dogs did not experience particular contexts while at the shelter. The missing data rate was between 0.06% and 5% for each context, except for the Interactions with female dogs and Interactions with male dogs categories which had 17% and 18% of missing values, respectively (because structured interactions with other dogs did not arise as frequently). Moreover, 16% and 8% of dogs were missing weight measurement and neuter status data, respectively, which were independent variables statistically controlled for in subsequent analyses. We created 5 multiply imputed data sets (using the *Amelia* package; [61]), upon which all following analyses in the sections below were conducted and results pooled. The multiple imputation took into account the hierarchical structure of the data (observations within dogs), all independent variables reported below, and the data types (ordered binary variables for the context data, positive-continuous for weight measurements, nominal for neuter status; see the R script). The data were assumed to be missing at random, that is, dependent only on other variables in the analyses.

### Structural equation models

We used structural equation modelling to assess whether aggression towards people(contexts: Handling, In kennel towards people, Out of kennel towards people, Interactions with familiar people, Interactions with unfamiliar people, Eating food, Interactions with toys) and towards dogs (contexts: In kennel towards dogs, Out of kennel towards dogs, Interactions with female dogs, Interactions with male dogs) could be explained by two latent aggressiveness traits: aggressiveness towards people and dogs, respectively. Since positive correlations between different aggressiveness traits have been reported in dogs [55], we compared a model where the latent variables were orthogonal to a model where variables were allowed to covary. Models were fit using the *lavaan* package [62], with the weighted least squares mean and variance adjusted (WLSMV) estimator and theta/conditional parameterisation, as recommended for categorical dependent variables [8,63,64]. The latent variables were standardised to have mean 0 and variance 1. The results were combined across imputed data sets using the ‘runMI’function in the *semTools* package [65]. The fit of each model was ascertained using the comparative fit index (CFI) and Tucker Lewis index (TLI), where values > 0.95 indicate excellent fit, as well as the root mean squared error of approximation (RMSEA) where values < 0.06 indicate good fit [7]. Parameter estimates were summarised by test statistics and 95% confidence intervals (CI).

### Local independence

We tested the assumption of local independence by re-fitting the best-fitting structural equation model with residual covariances specified between context variables. To maintain a theoretically driven approach (see [66] regarding the best practice of including residual covariances in structural equation models) and model identifiability, we only tested a predefined set of covariances based on which contexts shared close temporal-spatial relationships. First, we allowed covariances between *Handling* with *In kennel* towards people, Interactions with familiar people, Interactions with unfamiliar people and Interactions with toys, respectively, since the Handling context could directly succeed these other contexts. The residual covariance between Handling and Eating food was not estimated because shelter employees would be unlikely to handle a dog while the dog ate its daily meals. The residual covariance between *Handling* and *Out of kennel towards people* was not estimated because any association between *Handling* and *Out of*kennel towards people would be mediated by either the Interactions with familiar people or Interactions with unfamiliar people context. Therefore, secondly, we estimated the three-way covariances between Out of kennel towards people, Interactions with familiar people and Interactions with unfamiliar people. Similarly, and lastly, we estimated the three-way covariances between Out of kennel towards dogs, Interactions with female dogs and Interactions with male dogs. No covariances were inspected between In kennel towards dogs and other aggressiveness towards dogs contexts since large time gaps were more likely to separate observations between those contexts.

### Measurement invariance

To test for measurement invariance in each of the latent traits derived from the best fitting structural equation model, we investigated the response patterns across aggression contexts related to the same latent aggressiveness trait using Bayesian hierarchical logistic regression models. These models were analogous to the 1-parameter item response theory model, which represents the probability that an individual responds correctly to a particular test item as a logistic function of i) each individual’s latent ability and ii) the item’s difficulty level. This model can be expressed as a hierarchical logistic regression model [67,68], whereby individual latent abilities are modelled as individual-specific intercepts (i.e. ‘random intercepts’), the propensity for a correct answer to an item *i* is its regression coefficient β_i_, and credible interactions between items and relevant independent variables (e.g. group status) indicate a violation of measurement invariance. Here, the dependent variable was the binary score for whether or not dogs had shown aggression in each context and the average probability of aggression across contexts varied by dog, representing latent levels of aggressiveness. Context type, dog age, dog sex and their interactions were included as categorical independent variables. Age was treated as a categorical variable, with categories reflecting general developmental periods: i) 4 months to 10 months (juvenile dogs before puberty), ii) 10 months to 3 years (dogs maturing from juveniles to adults), iii) 3 years to 6 years (adults), and iv) 6 years + (older dogs). Broad age categories were chosen due to potentially large differences in developmental timing between individuals. Age was categorised because we predicted that aggression would be dependent on these developmental periods.

Models included additional demographic variables (Table 1) that may mediate the probability of aggression: body weight (average weight if multiple measurements were taken), total number of days spent at the shelter, the rehoming centre at which dogs were based (London, Old Windsor, Brands Hatch), neuter status (neutered before arrival, neutered at the shelter, not neutered) and source type (relinquished by owner, returned to the shelter after adoption, stray). Categorical variables were represented as sum-to-zero deflections from the group-level intercept to ensure that the intercept represented the average probability of aggression across the levels of each categorical predictor. Weight and total days at the shelter were mean-centered and standardised by 2 standard deviations. Due to the potentially complex relationships between these variables and aggression (e.g. interactive effects between neuter status and sex; [52]), which could also include violations of measurement invariance, we decided not to interpret their effects inferentially. Instead, they were included to make the assessment of measurement invariance between sexes and age groups conditional on variance explained by potentially important factors.

For comparability to other studies in animal personality, behavioural repeatability was calculated across contexts in each model using the intraclass correlation coefficient(ICC), calculated as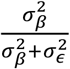, where 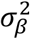represented the between-individual variance of the probability of aggression (i.e. the variance of the random intercepts), and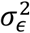, the residual variance of the standard logistic distribution [69].

#### Computation

Models were computed using the probabilistic programming language Stan version 2.15.1 [70], using Hamiltonian Monte Carlo, a type of Markov Chain Monte Carlo (MCMC) algorithm, to sample from the posterior distribution. Prior distributions for all independent variables were normal distributions with mean 0 and standard deviation 1, attenuating regression coefficients towards zero for conservative inference. The prior on the overall intercept parameter was normally distributed with mean 0 and standard deviation 5. The standard deviation of dog-specific intercept parameters was given a half-Cauchy prior distribution with mean 0 and shape 2. Each model was run with 4 chains of 2,000 iterations with a 1,000 step warm-up period. The Gelman-Rubin statistic (ideally < 1.05) and visual assessment of traceplots were used to assess MCMC convergence. We checked the accuracy of the model predictions against the raw data using graphical posterior predictive checks. For plotting purposes, predicted probabilities of aggression were obtained by marginalising over the random effects (explained in the Supporting Information). Regression coefficients were expressed as odds ratios and were summarised by their mean and 95% Bayesian highest density interval (HDI), representing the 95% most probable parameter values. To compare levels of categorical variables and their interactions, we computed the 95% HDI of the differences between the respective posterior distributions.

#### Model selection & parameter inference

Models were run on each imputed data set and their respective posterior distributions were averaged to attain a single posterior distribution for inference. Adopting a Bayesian approach allowed the estimation of interaction parameters (i.e. testing measurement invariance) without requiring corrections for multiple comparisons as in null hypothesis significance testing [71]. Nonetheless, models included a large number of estimated parameters. Two strategies were employed to guard against over-fitting of models to data. First, we selected the model with the best out-of-sample predictive accuracy given the number of parameters based on the Widely Applicable Information Criterion (WAIC; using the R package *loo* [72]). Four variants of each model were computed: two-way interactions between contexts and age and contexts and sex, respectively (model 1), a single interaction with sex but not with age (model 2), a single interaction with age but not with sex (model 3), and no interactions (model 4). All models included the mediating independent variables above. Second, to avoid testing point-estimate null hypotheses, the effect of a parameter was only considered credibly different from zero if the odds ratio exceeded the region of practical equivalence (ROPE; see [73]) around an odds ratio of 1 from 0.80 to 1.25. An odds ratio of 0.80 or 1.25 indicates a 20% decrease or increase (i.e. 4/5 or 5/4 odds), respectively, in the odds of an outcome, frequently used in areas of bioequivalence testing (e.g. [74]), which we here considered to be small enough to demonstrate a negligible effect in the absence of additional information. If a 95% HDI fell completely within the ROPE, the null hypothesis of no credible influence of that parameter was accepted; if a 95% HDI included part of the ROPE, then the parameter’s influence was left undecided [73].

### Ethics statement

Permission to use and publish the data was received from the shelter. Approval from an ethical review board was not required for this study.

### Data accessibility

Supporting Information (data, R script, Stan model code, Tables S1–4) can be found at:https://github.com/ConorGoold/GooldNewberry_aggression_shelter_dogs.

## Results

### Validity of behaviour recordings

For the video showing aggression towards people, 52% of participants identified the behaviour correctly as aggression and 42% identified the behaviour as non-aggressive but (similarly) reactive behaviour (see definitions above). For the video showing aggression towards dogs, 53% identified the behaviour correctly and 44% identified the behaviour as non-aggressive but reactive behaviour. For the 12 other videos not showing aggression, only 1 person incorrectly coded a video as aggression towards people and 3 people incorrectly coded videos as aggression towards dogs.

### Structural equation models

The raw tetrachoric correlations between the aggression contexts were all positive, particularly between contexts recording aggression towards people and dogs, respectively, supporting their convergent validity (Table S2). The model with correlated latent variables fit marginally better (CFI: 0.96; TLI: 0.95; RMSEA: 0.03) than the model with uncorrelated variables (CFI: 0.94; TLI: 0.92; RMSEA: 0.04). All regression coefficients of the model with correlated latent variables were positive and significant (i.e. the 95% CI did not include zero), and the latent variables shared a significant positive covariance (Table 3).

**Table 3.** Parameter estimates from the best-fitting structural equation model.

| Parameter | Estimate | SE | *t* value | 95% CI |
| --- | --- | --- | --- | --- |
| Handling<sup>a</sup> | 0.81 | 0.06 | 14.25 | [0.70, 0.92] |
| In kennel towards people<sup>a</sup> | 1.29 | 0.09 | 14.17 | [1.12, 1.46] |
| Out of kennel towards people<sup>a</sup> | 0.83 | 0.07 | 11.99 | [0.69, 0.96] |
| Interactions with familiar people<sup>a</sup> | 0.96 | 0.07 | 14.23 | [0.83, 1.09] |
| Interactions with unfamiliar people<sup>a</sup> | 1.54 | 0.12 | 12.46 | [1.23, 1.78] |
| Eating food<sup>a</sup> | 0.70 | 0.06 | 12.33 | [0.59, 0.81] |
| Interactions with toys<sup>a</sup> | 0.51 | 0.06 | 8.32 | [0.39, 0.63] |
| In kennel towards dogs<sup>b</sup> | 0.70 | 0.06 | 11.94 | [0.59, 0.82] |
| Out of kennel towards dogs<sup>b</sup> | 0.47 | 0.04 | 10.80 | [0.38, 0.55] |
| Interactions with female dogs<sup>b</sup> | 0.87 | 0.07 | 12.05 | [0.72, 1.02] |
| Interactions with male dogs<sup>b</sup> | 0.88 | 0.07 | 12.23 | [0.74, 1.03] |
| Covariance: People ~ Dogs | 0.26 | 0.03 | 7.94 | [0.19, 0.33] |

<sup>a</sup> Contexts reflecting aggressiveness towards people
<sup>b</sup> Contexts reflecting aggressiveness towards dogs

### Local independence

Allowing the pre-defined residuals to co-vary in the best-fitting structural equation model resulted in a better fit (CFI = 0.98; TLI = 0.97; RMSEA: 0.03). Significant negative covariances were observed between the *Handling* and *In kennel towards people* contexts (Table 4) and the *Handling* and *Interactions with unfamiliar people* contexts. A significant positive covariance was observed between *Out of kennel towards people* and *Interactions with unfamiliar people* contexts. No significant residual covariances between contexts reflecting aggressiveness towards dogs were observed.

**Table 4.**
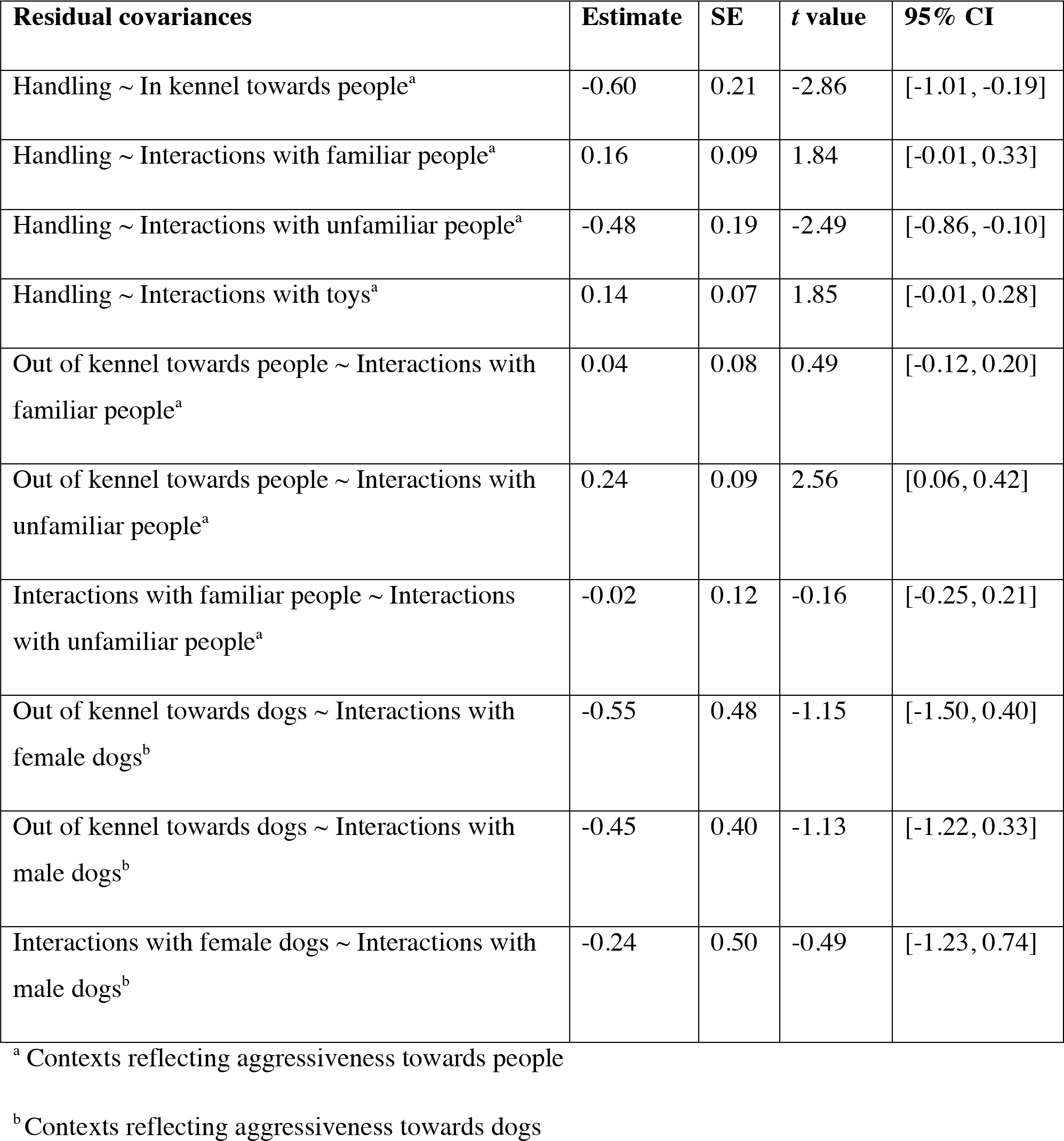
Estimated residual covariances between contexts.

| **Residual covariances** | **Estimate** | **SE** | ***t* value** | **95% CI** |
| --- | --- | --- | --- | --- |
| Handling ~ In kennel towards people<sup>a</sup> | -0.60 | 0.21 | -2.86 | [-1.01, -0.19] |
| Handling ~ Interactions with familiar people<sup>a</sup> | 0.16 | 0.09 | 1.84 | [-0.01, 0.33] |
| Handling ~ Interactions with unfamiliar people<sup>a</sup> | -0.48 | 0.19 | -2.49 | [-0.86, -0.10] |
| Handling ~ Interactions with toys<sup>a</sup> | 0.14 | 0.07 | 1.85 | [-0.01, 0.28] |
| Out of kennel towards people ~ Interactions with familiar people<sup>a</sup> | 0.04 | 0.08 | 0.49 | [-0.12, 0.20] |
| Out of kennel towards people ~ Interactions with unfamiliar people<sup>a</sup> | 0.24 | 0.09 | 2.56 | [0.06, 0.42] |
| Interactions with familiar people ~ Interactions with unfamiliar people<sup>a</sup> | -0.02 | 0.12 | -0.16 | [-0.25, 0.21] |
| Out of kennel towards dogs ~ Interactions with female dogs<sup>b</sup> | -0.55 | 0.48 | -1.15 | [-1.50, 0.40] |
| Out of kennel towards dogs ~ Interactions with male dogs<sup>b</sup> | -0.45 | 0.40 | -1.13 | [-1.22, 0.33] |
| Interactions with female dogs ~ Interactions with male dogs<sup>b</sup> | -0.24 | 0.50 | -0.49 | [-1.23, 0.74] |

<sup>a</sup> Contexts reflecting aggressiveness towards people
<sup>b</sup> Contexts reflecting aggressiveness towards dogs

### Measurement invariance

Separate models were run for contexts reflecting aggressiveness towards people and aggressiveness towards dogs. All models converged. Posterior predictive checks of model estimates reflected the raw data (Figs 1 and 2). The full measurement invariance model (model 1) including interactions between contexts and sex and contexts and age groups had the best out-of-sample predictive accuracy for both the aggressiveness towards people and aggressiveness towards dogs models, respectively, illustrated by the lowest WAIC values (Table 5). Since some models included numerous interactions, we provide an overall summary of the main results below (Figs 1 and 2) with full parameter estimates provided in Tables S3 and S4.

**Table 5.** Mean ± standard error of the Widely Applicable Information Criteria (WAIC) values (lower is better) per model and aggressiveness variable.

| Model | Aggressiveness towards people | Aggressiveness towards dogs |
| --- | --- | --- |
| Model 1 | 13405.6 ± 179.0 | 15257.2 ± 133.1 |
| Model 2 | 13506.3 ± 179.6 | 15381.4 ± 133.4 |
| Model 3 | 13426.3 ± 179.1 | 15285.3 ± 133.0 |
| Model 4 | 13521.7 ± 179.5 | 15407.6 ± 133.4 |

#### Aggressiveness towards people

The odds of aggression towards people, across categorical predictors and for an average dog of mean weight and length of stay at the shelter, were 0.022 (HDI: 0.021 to 0.024), a probability of approximately 2%. On average, aggression was most likely in the *In kennel towards people* context (OR = 0.054; HDI: 0.049 to 0.058) and least probable in the*Interactions with toys* context (OR = 0.008; HDI: 0.007 to 0.009).

Aggression was less likely across contexts for females than males (OR = 0.719; HDI: 0.668 to 0.770), although there were also credible interactions between sex and contexts (Fig 1A; Table S3). Whereas males and females had similar odds of aggression in the *Out of kennel towards people* context, smaller differences were observed between *Out of kennel towards people* and *Handling* (OR = 0.578; HDI: 0.481 to 0.682), *Eating food* (OR = 1.812; HDI: 1.495 to 2.152) and *Interactions with familiar people* (OR = 1.798; HDI: 1.488 to 2.126) contexts in females compared to males. Additionally, whereas aggression in the *Interactions with unfamiliar people* context was similar between males and females, larger differences were observed between *Interactions with unfamiliar people* and *Handling* (OR = 0.616; HDI: 0.530 to 0.702), *Eating food* (OR = 0.594; HDI:0.506 to 0.686) and *Interactions with familiar people* (OR = 0.598; HDI: 0.513 to 0.687) contexts in females compared to males.

Apart from lower odds of aggression in 4 to 10 month olds compared to 10 month to 3 year old dogs (OR = 0.638; HDI: 0.565 to 0.705), there was no simple influence of age group on aggressiveness. Between the 4 to 10 months old and 3 to 6 years old groups, differences between the odds of aggression across contexts varied due to an increase of aggression in certain contexts but not others (Fig 1B; Table S4). Aggression in *In kennel towards people* and *Interactions with unfamiliar people* contexts particularly increased, leading to larger differences between, for example, *In kennel towards people* and *Eating food* (OR = 0.524; HDI: 0.400 to 0.642) and *Eating food* and *Interactions with unfamiliar people* (OR = 1.721; HDI: 1.403 to 2.059) contexts for 10 month to 3 year olds compared to 4 to 10 month olds, and between *In kennel towards people* and *Out of kennel towards people* (OR = 0.470; HDI: 0.355 to 0.606) and *Out of kennel towards people* and *Interactions with unfamiliar people* (OR = 2.051; HDI: 1.608 to 2.543) contexts in 3 to 6 year olds compared to 4 to 10 month olds. In 3 to 6 year old compared to 10 month to 3 year old dogs, aggression increased in the *Handling* and *Eating food* contexts but decreased in the *Out of kennel towards people* context, resulting in larger differences between, for instance, *Handling* and *Out of kennel towards people* (OR = 0.526; HDI:0.409 to 0.631) and *Out of kennel towards people* and *Interactions with unfamiliar people* (OR = 2.349; HDI: 1.891 to 2.925), and smaller differences between *Eating food* and *Interactions with familiar people* (OR = 0.576; HDI: 0.468 to 0.687).

Dogs over 6 years old demonstrated qualitatively different response patterns across certain contexts than all other age groups. While aggression was most probable in *In kennel towards people* and *Interactions with unfamiliar people* contexts for dogs aged 4 months through 6 years, dogs over 6 years old were most likely to show aggression in the *Handling* context, leading to interactions between, for example, *Handling* and *In kennel towards people*, and between *Handling* and *Interactions with unfamiliar people* contexts compared to the other age groups (Fig 1B; Table S3). Aggression when *Eating food* and in *Interactions with toys* contexts also increased compared to that expressed by younger dogs, resulting in credible differences between, for instance, *Eating food* and *Interactions with familiar people* contexts between dogs aged 10 months to 3 years and over 6 years (OR = 0.379; HDI: 0.300 to 0.465) and between *Out of kennel towards people* and*Interactions with toys* contexts between over 6 year olds and all other age groups (Table S3).

**Figure 1.**
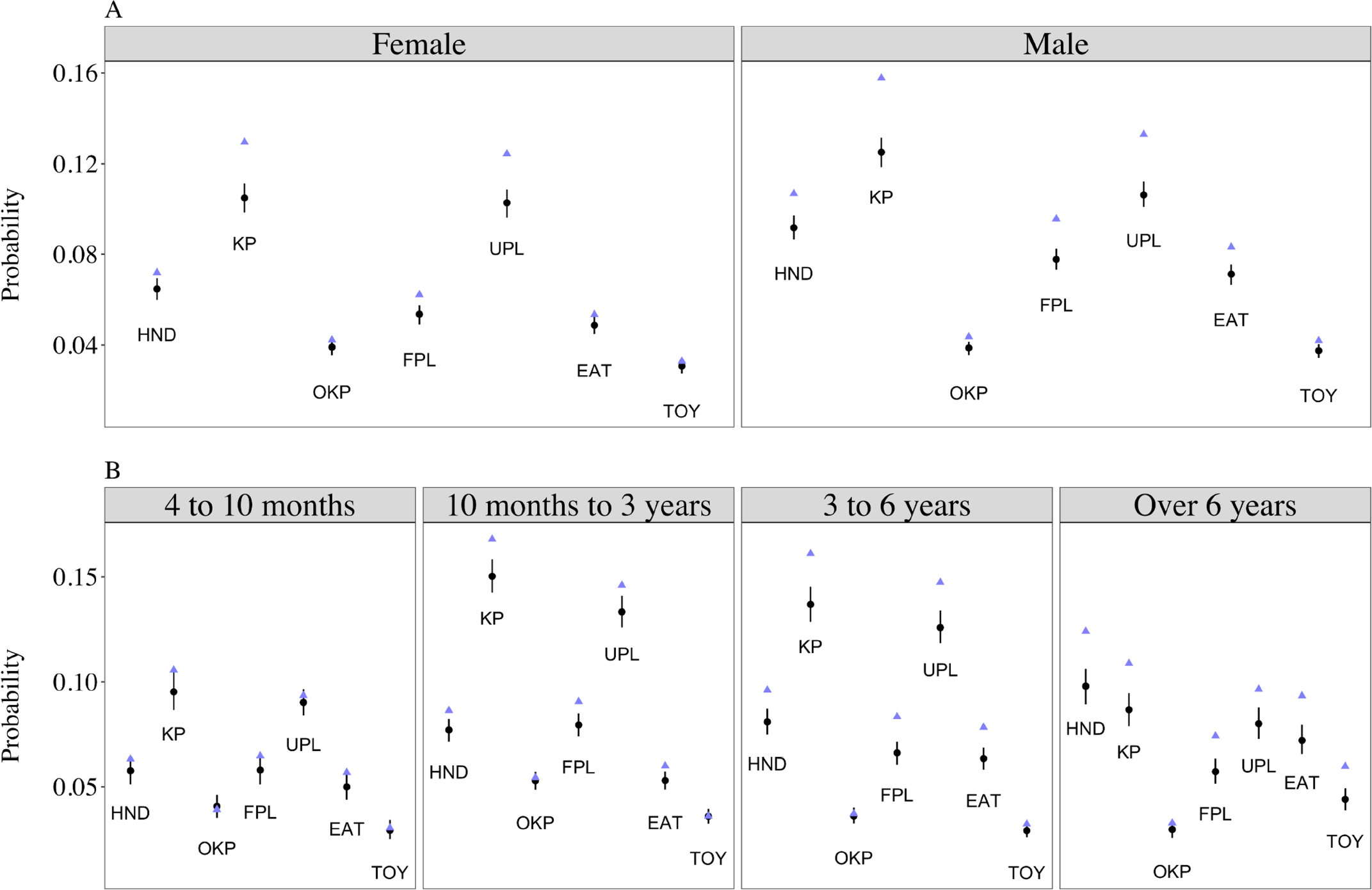
Predicted probabilities of aggression towards people in different contexts by sex (panel A) and age groups (panel B). Black points and vertical lines show mean and 95% highest density intervals of model parameter estimates; blue triangles show raw sample data. Model estimates were obtained by marginalising over the random effects(see the Supporting Information). Abbreviations used in the figure: HND (*Handling*); KP (*In kennel towards people*); OKP (*Out of kennel towards people*); FPL (*Interactions with familiar people*); UPL (*Interactions with unfamiliar people*); EAT (*Eating food*); TOY (*Interactions with toys*).

#### Aggressiveness towards dogs

The odds of aggression towards dogs, across categorical predictors and for an average dog of mean weight and length of stay at the shelter, was 0.176 (HDI: 0.168 to 0.184), corresponding to a probability of approximately 15%. Dogs were most likely to show aggression in the *Interactions with male dogs* context (OR = 0.297; HDI: 0.198 to 0.217) and least likely in the *In kennel towards dogs* context (OR = 0.099; HDI: 0.094 to 0.104; Fig 2; Table S4).

No credible mean-level differences existed between females and males (OR = 1.187; HDI: 1.128 to 1.250). However, the difference in aggression between the *Interactions with female dogs* and *Interactions with male dogs* contexts was smaller for females (OR = 1.542; HDI: 1.400 to 1.704; Fig 2A; Table S4), as were the differences between *Interactions with male dogs* and *In kennel towards dogs* (OR = 0.661; HDI: 0.590 to 0.732) and *In kennel towards dogs* and *Out of kennel towards dogs* (OR = 1.420; HDI:1.269 to 1.587). Females were also more likely to show aggression in *Interactions with female dogs* than *Out of kennel towards dogs* compared to males (OR = 1.444; HDI: 1.301 to 1.603).

Dogs aged 4 to 10 months old had credibly lower odds of aggression towards dogs than older dogs across contexts (Fig 2B; Table S4). However, contexts and age also showed interactive effects. In particular, aggression in *Interactions with female dogs* and*Interactions with male dogs* contexts tended to increase relative to other contexts. For instance, the relationship between *Interactions with female dogs* and *Out of kennel towards dogs* contexts reversed in direction between 4 to 10 month and 10 month to 3 year olds (OR = 0.595; HDI: 0.495 to 0.688) as did the relationship between *Interactions with male dogs* and *Out of kennel towards dogs* contexts (OR = 0.499; HDI: 0.422 to 0.575). The relationship between *In kennel towards dogs* and *Out of kennel towards dogs*contexts also changed across age groups (Fig 2B; Table S4). Four to 10 months old were more likely to show aggression in *Out of kennel towards dogs* than *In kennel towards dogs* contexts, but the difference was smaller in 10 months to 3 year olds (OR = 0.608; HDI: 0.505 to 0.728) and in over 6 year olds (OR = 0.396; HDI: 0.316 to 0.481). The latter relationship was reversed in 3 to 6 year olds compared to 4 to 10 month old dogs (OR = 0.277; HDI: 0.227 to 0.331) and 10 month to 3 year old dogs (OR = 0.456; HDI:0.396 to 0.516).

**Figure 2.**
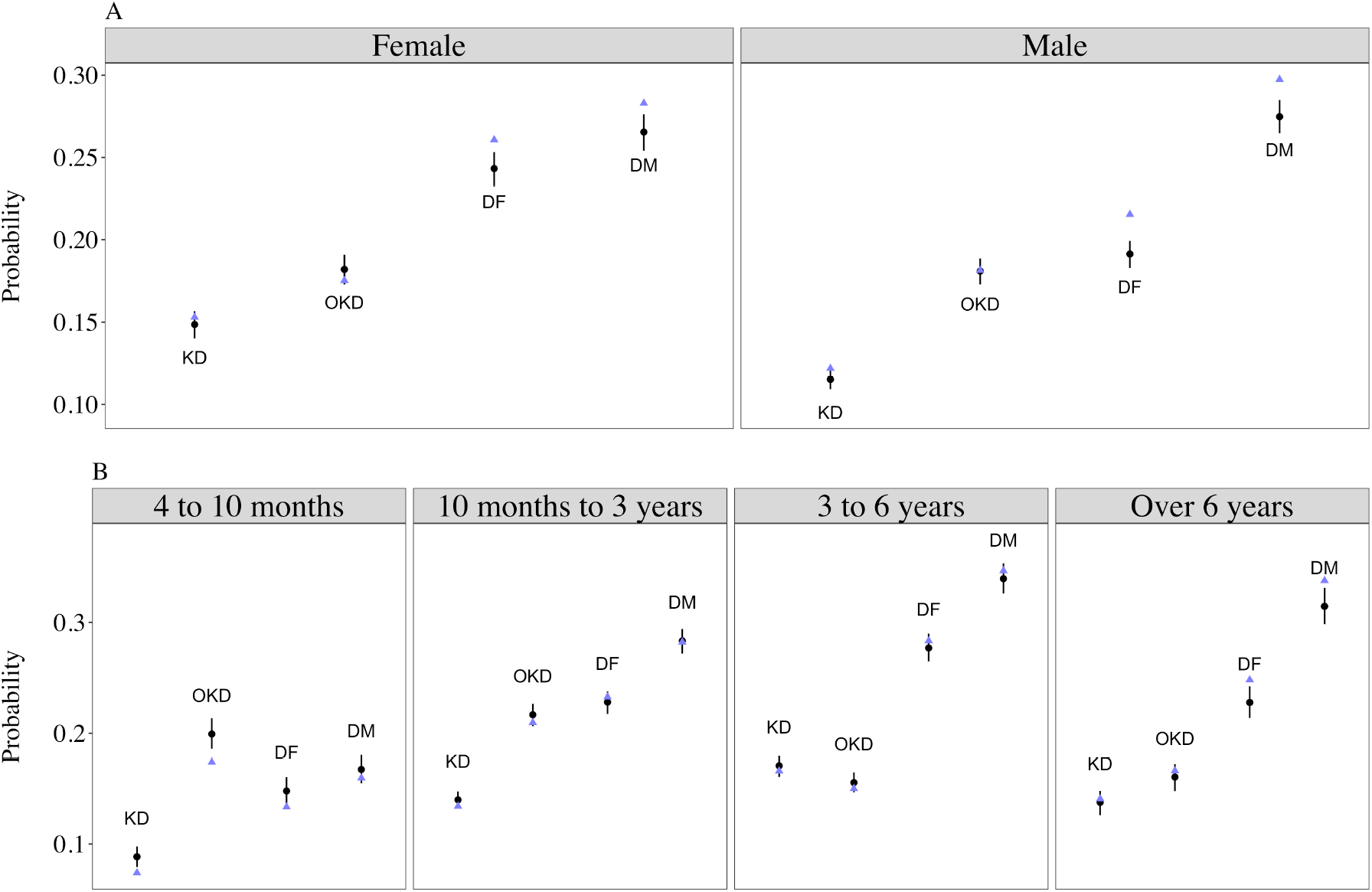
**Predicted probabilities of aggression towards dogs in different contexts by sex (panel A) and age groups (panel B)**. Black points and vertical lines show mean and 95% highest density intervals of model parameter estimates; blue triangles show raw sample data. Model estimates were obtained by marginalising over the random effects (see the Supporting Information). Abbreviations used in the figure: KD (*In kennel towards dogs*); OKD (*Out of kennel towards dogs*); DF (*Interactions with female dogs*); DM (Interactions with male dogs).

## Repeatability

Both aggressiveness towards people and dogs showed moderate repeatability across contexts (*ICC_people_*= 0.479; HDI: 0.466 to 0.491; *ICC_dogs_* = 0.303; HDI: 0.291 to 0.315), although aggressiveness towards people was more repeatable than aggressiveness towards dogs (*ICC_difference_*= 0.176; HDI: 0.158 to 0.192).

## Discussion

In this study, we have examined local independence and measurement invariance of aggressiveness traits in shelter dogs. Observational recordings of aggression directed towards people and dogs across different shelter contexts were explained by two positively correlated latent variables, and behaviour across contexts was moderately repeatable. These results are consistent with the concept of animal personality, which is used to describe behaviour that shows moderately consistent between-individual differences across time or contexts, and is characterised by multiple observed behaviours being decomposed into lower-dimensional behavioural traits [4]. However, we found violations of local independence between contexts with close temporal-spatial relationships and measurement invariance with respect to sex and age groups, highlighting potential measurement biases.

Local independence implies that the association between manifest variables is greater than that explained by the latent variable. For aggressiveness towards people, aggression in the *Handling* context was negatively related with the *In kennel towards people* and *Interactions with unfamiliar people* contexts, while positive covariances were present between Out of kennel towards people and Interactions with unfamiliar people contexts. Violations of local independence may arise through shared method variance [75–78] or unmodelled latent variables influencing manifest variables [79,80]. If a dog showed aggression when an unfamiliar person approached, it may be less likely to be handled by that person, which may explain the negative residual covariations between the *Handling* and *In kennel towards people* and *Interactions with unfamiliar people* contexts, respectively. These contexts were, in fact, positively correlated when latent levels of aggressiveness were not accounted for (Table S4). In addition, the positive residual correlation between *Out of kennel towards people* and *Interactions with unfamiliar people* may be mediated by additional traits of interest to personality researchers, such as fearfulness or anxiety [29,81], if dogs who are fearful of interacting with unfamiliar people are more likely to show aggression beyond that described by a latent aggressiveness trait.

While authors have argued that greater standardisation and validation of personality assessments is key to ensuring the accurate measurement of underlying traits [36,48,49], it may be untenable to avoid dependencies between testing contexts. Displays of aggression in one sub-test will likely change how people conduct future sub-tests with the same dog, regardless of test standardisation. Human psychologists have argued that violations of local independence are a natural consequence of the organisation of behaviour as a complex dynamic system [82,83], which unfolds with respect to time- and context-dependent constraints [84]. Thus, awareness of local independence and its violation could facilitate closer understanding of the dynamics driving personality test responses beyond explanations purely based on personality traits.

While different subsets of a population may differ in mean levels of trait expression, interactions between behavioural responses and those subsets indicate that the same phenomenon is not under measurement across groups [23,24]. We found that the probability of aggression across contexts was dependent on sex and age conditional on latent levels of aggressiveness (Figs 1 and 2; Tables S3 and S4). Female dogs, for example, were more likely than males to show aggression in *Out of kennel towards people* and *Interactions with unfamiliar people* contexts relative to other contexts (Fig 1A). Females also demonstrated similar odds of aggression during *Interactions with female dogs* and *Interactions with male dogs*, whereas males were more likely to show aggression towards male than female dogs (Fig 2a). As with local independence, different behavioural variables unaccounted for in this study may result in violations of measurement invariance. While dogs up to 6 years old were most likely to show aggression in *In kennel towards people* and *Interactions with unfamiliar people contexts*, dogs over 6 years old demonstrated aggression most commonly in the *Handling* context. Dogs over 6 years old also showed an increase in aggression in the *Eating food* and *Interactions with toys* contexts relative to other age groups. These results suggest that older dogs in shelter populations may be less tolerant during close interactions with people (i.e. handling, people in the vicinity of their food and toys) compared to other contexts, which may driven by other quantifiable factors such as pain or sensitivity (e.g. [29]).

Although we have identified violations of both local independence and measurement invariance, we remain cautious about hypothesising *a posteriori* about their causes. Personality traits in animal behaviour are typically defined operationally, based on the statistical repeatability of quantifiable behaviour [77,85,86]. As discussed in human personality psychology, operational definitions can be ontologically ambiguous [87,88]. That is, while operational definitions facilitate experimentation in animal personality [4], they do not necessarily designate biological mechanisms underlying trait expression. For example, Budaev and Brown remark that boldness, defined as a propensity to take risks, could encompass a range of distinct personality traits, each with a different biological basis [75]. Whilst reflective latent variable models allow researchers to test hypotheses about the relatedness of measured behaviours via one or more underlying traits, they have also been criticised as ambiguous [82]. For example, it is uncertain what reflective latent variables may represent in biological organisation [87] or even whether they are features individuals possess or simply emergent features of between-individual differences [89,90]. Such considerations highlight the importance of research on the proximate mechanisms of personality [85] and longitudinal data analyses to separate between- from within-individual behavioural variation [91,92].

A number of authors have emphasised the poor predictive value of aggression tests in shelter dogs [39–41,50] and that low occurrence of aggression specifically can make its accurate measurement difficult [40]. The probability of observing aggression on any particular day was low in this study (approximately 1%), and the number of dogs who, on average, showed aggression to people at least once while at the shelter was much lower than the number that showed aggression towards dogs, on average (Figs 1 and 2). Nonetheless, our evaluations of validity indicated that between 40 and 45% of the shelter employees mistook observations of aggression for non-aggressive responses (e.g. over-excitement and frustration when seeing people/dogs), meaning that the true probability of aggression was potentially under-estimated (although incorrectly coding other behaviours as aggression also occurred, albeit rarely). Moreover, our assessments of validity were based on shelter staff evaluations of brief video recordings that may be less reliable than the live, spontaneous behavioural recordings upon which our main analyses were based, resulting in a lower percentage of correctly identified instances of aggression. For the two videos being evaluated, the shelter employees had 13 and 11 different behavioural codes, respectively, to choose from to describe the behaviours observed. Thus, while employees as a whole were undecided about whether the motivation for the behaviour was aggressive or non-aggressive, the vast majority of employees described the behaviour as reactive, despite potentially erring on the side of caution by labelling aggressive behaviours as non-aggressive. Comparable estimates of validity are not present in the literature on dog personality, but are particularly important in shelter settings where accurate recording of aggression is paramount. It is also worth noting that how to assess validity has received much debate (e.g. [87,92]). In this study, we used expert judgement as a benchmark to which shelter employees’ responses were compared, but validity is frequently assessed in dog personality by inspecting patterns of correlation coefficients between similar and dissimilar behaviours (e.g. convergent or divergent validity; [29]). This is less directly interpretable than reporting the percentage of answers that were correct, as used here. Moreover, the predictive validity of personality assessments in dogs have been inconsistent (e.g. [40–42]). More discussion of the concept of validity, and how best to assess it, is warranted in studies of dog personality.

Infrequent occurrence and/or recording of aggression may also limit accurate predictions of future behaviour. Patronek and Bradley [50] demonstrate using simulation that the low prevalence of aggression inflates the chance that aggression shown in a shelter assessment represents a false positive. In general, our results support this conclusion in the sense that aggression may be shown differentially across contexts not explained by latent levels of aggressiveness. Violations of local independence and measurement invariance as found here indicate, further, that it is not only the difference between false and true positives and negatives, but the validity of inferring homogeneous personality traits by which to compare individual dogs, that needs careful consideration.

Consequently, we agree with recommendations to establish the efficacy of longitudinal, observational assessments rather than relying on a single assessment made using a traditional test battery [31,40,50]. This approach will prioritise the cumulative understanding of a dog’s context-dependent behaviour and help to guide decisions about the potential risk a dog poses to humans and other animals.

## Conclusion

This study has tested the assumptions of local independence and measurement invariance of personality traits in shelter dogs. Using structural equation modelling, aggression across behavioural contexts was explained by two correlated latent variables and demonstrated repeatability. Nevertheless, significant residual covariances remained between certain behavioural contexts related to aggressiveness towards people, violating the assumption of local independence. In addition, aggression in different contexts showed differential patterns of response across sex and age, indicating a lack of measurement invariance. Violations of local independence and measurement invariance imply that the aggressiveness towards people and dogs traits did not completely explain patterns of aggression in different contexts, or that inferences based on these hypothesised personality traits may in fact be misleading. We encourage researchers to more closely assess the measurement assumptions underlying reflective latent variable models before making conclusions about the effects of, or factors influencing, personality.

## Acknowledgements

The authors are extremely grateful to Battersea Dogs and Cats Home for allowing us to access the data for this study.

---

**Table S1.** Counts of aggression per context. The number of dogs who had 0, 1, and > 1 observations of aggression while at the shelter.

**Table S2.** Tetrachoric correlations between aggression contexts. Tetrachoric correlations between aggression contexts on the raw binary data, before the multiple imputation. Abbreviations used: HND (*Handling*); FPL (*Interactions with familiar people*); UPL (*Interactions with unfamiliar people*); KD (*In kennel towards dogs*); KP (*In kennel towards people*); OKD (*Out of kennel towards dogs*); OKP (*Out of kennel towards people*); EAT (*Eating food*); TOY (*Interactions with toys*); DM (*Interactions with male dogs*); DF (*Interactions with female dogs*).

**Table S3.** Bayesian hierarchical model parameter estimates for aggression towards people in different contexts. Mean and 95% highest density interval (HDI) estimates for all parameters from the Bayesian hierarchical logistic model assessing measurement invariance for contexts reflecting aggressiveness towards people. Differences between levels of categorical variables are indicated by ‘.v.’ in the parameter name; interactions are denoted with ‘*’ in the parameter name. The decision rule for each parameter is given except for those variables not interpreted inferentially: YES = 95% HDI falls completely outside the region of practical equivalence (ROPE); NULL = 95% HDI falls completely inside the ROPE; ROPE = 95% HDI partly covers the ROPE.

**Table S4.** Bayesian hierarchical model parameter estimates for aggression towards dogs in different contexts. Mean and 95% highest density interval (HDI) estimates for all parameters from the Bayesian hierarchical logistic model assessing measurement invariance for contexts reflecting aggressiveness towards dogs. Differences between levels of categorical variables are indicated by ‘.v.’ in the parameter name; interactions are denoted with ‘*’ in the parameter name. The decision rule for each parameter is given except for those variables not interpreted inferentially: YES = 95% HDI falls completely outside the region of practical equivalence (ROPE); NULL = 95% HDI falls completely inside the ROPE; ROPE = 95% HDI partly covers the ROPE.

